# Memory-Phenotype Ly49+ CD8 T emerging from the thymus develop into two subsets with distinct immune functions

**DOI:** 10.1101/2024.12.10.627689

**Authors:** Daphné Laubreton, Morgan Grau, Victor Malassigne, Margaux Prieux, Martine Tomkowiak, Marc Vocanson, Thierry Walzer, Jacqueline Marvel

**Affiliations:** CIRI, Inserm U1111 - CNRS UMR5308 Universite Lyon1, ENS de Lyon

## Abstract

Ly49+ CD8 T cells are memory-phenotype (MP) cells expressing Ly49 family inhibitory receptors, that control autoimmune diseases development through the regulation of follicular CD4 T cells. During viral infection, their number increases, suggesting an involvement in antiviral responses. These cells do not derive from naive cells and are thought to develop in the thymus. This study identified two subsets of Ly49+ CD8 T cells, based on CD8β expression, in mice and humans. Lineage tracing and reliance on the transcription factor Zeb1 indicate a thymic origin via agonist selection. scRNAseq analysis revealed that a small fraction of CD8αβ-Ly49+ cells acquired an effector profile during vaccinia virus infection. Moreover, the majority of Ly49+ CD8 T cells seems to respond to cytokine-driven bystander signals. *In vitro*, these signals promoted the expression of effector molecules such as IFNγ and granzyme B, as well as the homing chemokine receptor CXCR5, potentially driving their recruitment to germinal centers.

**Summary:** This study shows that Ly49+ CD8 T cells comprises two subsets that develop in the thymus through agonist selection. In viral infection, both subsets respond to bystander stimuli that could drive their regulatory activity. Furthermore, some CD8αβ-expressing Ly49+ cells exert TCR-mediated antiviral response.

## Introduction

Following encounter with an intracellular pathogen or cancer cells, naive CD8 T cells are activated and differentiate in effector cells that eliminate infected or tumoral cells and memory cells that are responsible for host protection in case of antigenic rechallenge. However, some memory-phenotype (MP) CD8 T cells (also referred to as virtual-memory or innate memory cells) can be generated independently of antigen exposure (Thiele *et al*., 2020). These cells exhibit characteristics of antigen-experienced memory cells, such as CD44 and CD122 expression, and display enhanced responsiveness to antigenic but also cytokine stimuli (Thiele *et al*., 2020). Among MP CD8 T cells, there is a particular population referred to as Ly49+ CD8 T cells, that is characterized by the expression of Ly49A, Ly49F, Ly49G2, and, to a minor extent, Ly49C/I inhibitory receptors (Coles *et al*., 2000; Anfossi *et al*., 2004; Kim *et al*., 2011), and the expression of the EOMES (Mishra *et al*., 2021) and Helios (Kim *et al*., 2015) transcription factors.

Ly49+ CD8 cells play an important role in regulating germinal center reactions by controlling CD4 Tfh cells, thereby preventing the production of autoantibodies (Kim *et al*., 2010, 2015, 2024; Leavenworth *et al*., 2013; Alvarez Arias *et al*., 2014; Saligrama *et al*., 2019; Stocks *et al*., 2019; Mishra *et al*., 2021). The importance of the suppressive activity of Ly49+ CD8 T cells has been demonstrated in various pathologies, such as experimental autoimmune encephalitis (EAE) (Hu *et al*., 2004; Tang *et al*., 2006; Yu, Bamford and Waldmann, 2014; Saligrama *et al*., 2019), atherosclerosis (Clement *et al*., 2015), arthritis (Leavenworth *et al*., 2013), type 1 diabetes (Stocks *et al*., 2019) and colitis (Endharti *et al*., 2011; Yao *et al*., 2013). They can also prevent allograft rejection (Choi *et al*., 2020; Kim *et al*., 2024). The suppressive function of Ly49+ CD8 T cells appears to be TCR-mediated, requiring recognition of self-peptides presented by the non-classical MHC-I Qa-1 (Lu *et al*., 2008; Kim and Cantor, 2011) as well as classical-MHC-I (Li *et al*., 2022). Qa-1-D227K mutant mice, in which Qa-1:TCR binding is disrupted, spontaneously develop autoimmune disease symptoms, that are exacerbated following LCMV infection (Kim *et al*., 2010). Similarly, mice deficient for the Ly49F receptor, that have a profound defect in Ly49+ CD8 T cells, exhibit significant signs of autoimmunity within weeks following viral infection (Li *et al*., 2022). Interestingly, in Qa-1-D227K mutant mice, CD8 T cell responses to both acute and chronic LCMV infections (Holderried *et al*., 2013) or tumor challenge (Alvarez Arias *et al*., 2014) are enhanced, suggesting that Ly49+ CD8 T cells also regulate CD8 T cell responses via Qa-1. However, antiviral CD4 and CD8 T cell responses in Ly49F-KO mice are similar to those in WT mice (Li *et al*., 2022). This suggests that the regulation of CD8 T cells by Ly49+ CD8 T cells may rely on Qa-1-mediated signals, while the regulation of CD4 Tfh cells may involve both Qa-1 and classical MHC-I molecules. Finally, some studies report classical-MHC-I restricted responses by Ly49+ CD8 T cells to viral peptides (Peacock *et al*., 2000; Peacock and Welsh, 2004; Li *et al*., 2022). Overall, the antigenic peptide recognized and the stimuli driving the activation of these regulatory cells in a context of viral infection remain to be defined.

Some MP CD8 T cells are generated in the thymus in response to IL-4 produced by PLZF-expressing NKT cells (Jameson, Lee and Hogquist, 2015), while others are derived from naive cells in the periphery, in lymphopenic settings (Goldrath and Bevan, 1999; Kieper and Jameson, 1999) or following stimulation with γc-cytokines (Kamimura and Bevan, 2007; Ventre *et al*., 2012; Jason T. White *et al*., 2016). Unlike the latter, Ly49+ CD8 T cells cannot differentiate from naive CD8 T cells upon cytokine stimulation, whether in response to IL-2, IL-15, or both cytokines, even with the addition of TCR stimulation Similarly, naive CD8 T cells do not acquire Ly49 receptors expression after transfer into a Rag-/- host (Anfossi *et al*., 2004). Finally, the acquisition of Ly49 expression by naive CD8 T cells could also not be induced following diverse antigenic challenges, such as chronic or acute viral infection (Peacock *et al*., 2000; Peacock and Welsh, 2004). Altogether, these data suggest that Ly49+ CD8 T cells could thus be an independent subset of memory-phenotype CD8 T cells generated in the thymus, an hypothesis that is supported by the identification of CD8 single positive (SP) thymocytes expressing Ly49 receptors with a memory-phenotype (McCarron and Marie, 2014; Shytikov *et al*., 2021). However, the exact nature of Ly49+ CD8 T cells thymic precursors and the mechanisms driving their differentiation remain undefined.

In this study, we demonstrated that the expression of either the CD8αα or CD8αβ coreceptor defines two distinct subsets of Ly49+ CD8 T cells in mice as well as their human counterpart KIR+ CD8 T cells. Using multiparametric flow cytometry and pseudotime ordering, we mapped the developmental trajectory followed by Ly49+ CD8 T in the thymus. Our results suggest that these cells derive from a thymic precursor expressing EOMES, following an agonist selection process. In order to further characterize these subsets and their response during a viral infection we performed a scRNA-Seq transcriptomic analysis. This revealed that most Ly49+ CD8 T cells exhibit transcriptional changes indicative of a bystander response to viral infection, while a small fraction of CD8αβ-Ly49+ T cells is able to mount an antigen-specific effector response. Our findings provide a better comprehension of the origin and the functional role of Ly49+ CD8 T cells.

## Results

### CD8β expression defines two subsets of murine Ly49+ and human KIR+ CD8 T cells

The Ly49+ CD8 T cells described in the literature expressed mainly the CD8αβ coreceptor (Anfossi *et al*., 2004; Kim *et al*., 2011). However, one study reported the existence of Qa-1-restricted CD8αα-expressing T cells, that are enriched in the expression of Ly49A, Ly49F and Ly49G2 receptors (Fanchiang *et al*., 2012), suggesting that the Ly49+ CD8 T cell subset was heterogeneous. To address this question, we first monitored the expression of the CD8β chain by splenic Ly49+ CD8 T cells. Ly49+ CD8 T cells were selected according to their expression of at least one of the three inhibitory Ly49 receptors, Ly49A, Ly49F and Ly49G2 (Supplementary Fig. 1a). While all naive CD8 T cells express the CD8β chain, both CD8β+ and CD8β-cells were identified among Ly49+ CD8 T cells (Fig. 1a). In adult mice, CD8αα-Ly49+ cells represent 20 to 30% of the total pool of Ly49+ CD8 T cells (Fig. 1b). Next, we analyzed the Ly49 expression-pattern of CD8αα-Ly49+ and CD8αβ-Ly49+ cells. A large fraction of CD8αβ-Ly49+ cells express only one Ly49 receptor corresponding to Ly49F, with decreasing proportion of cells expressing 2 or 3 Ly49 receptors (Fig. 1c, Supplementary Fig. 1b). Conversely, CD8αα-Ly49+ cells have a similar representation of cells expressing 1, 2, or 3 different Ly49 receptors, with the Ly49F receptor being frequently associated with Ly49G2 alone or in combination with Ly49A (Fig. 1c, Supplementary Fig. 1b). The expression of the transcription factor Helios, one of the hallmark of Ly49+ CD8 T cells (Kim *et al*., 2015), was measured at the protein level. Its expression was observed in 80% of CD8αα-Ly49+ cells compared to less than 40% of CD8αβ-Ly49+ cells (Fig. 1d). Interestingly, within CD8αβ-Ly49+ cells but not CD8αα-Ly49+ cells, the expression of Helios correlated with the number of Ly49 receptors expressed (Fig. 1e). Thus, based on CD8β expression, we identified two Ly49+ CD8 cells subsets that differ in Helios expression. Furthermore, while both subsets can express the 3 Ly49 receptors (Ly49A, Ly49F and Ly49G2), they display different Ly49 expression-pattern.

**Figure 1:**
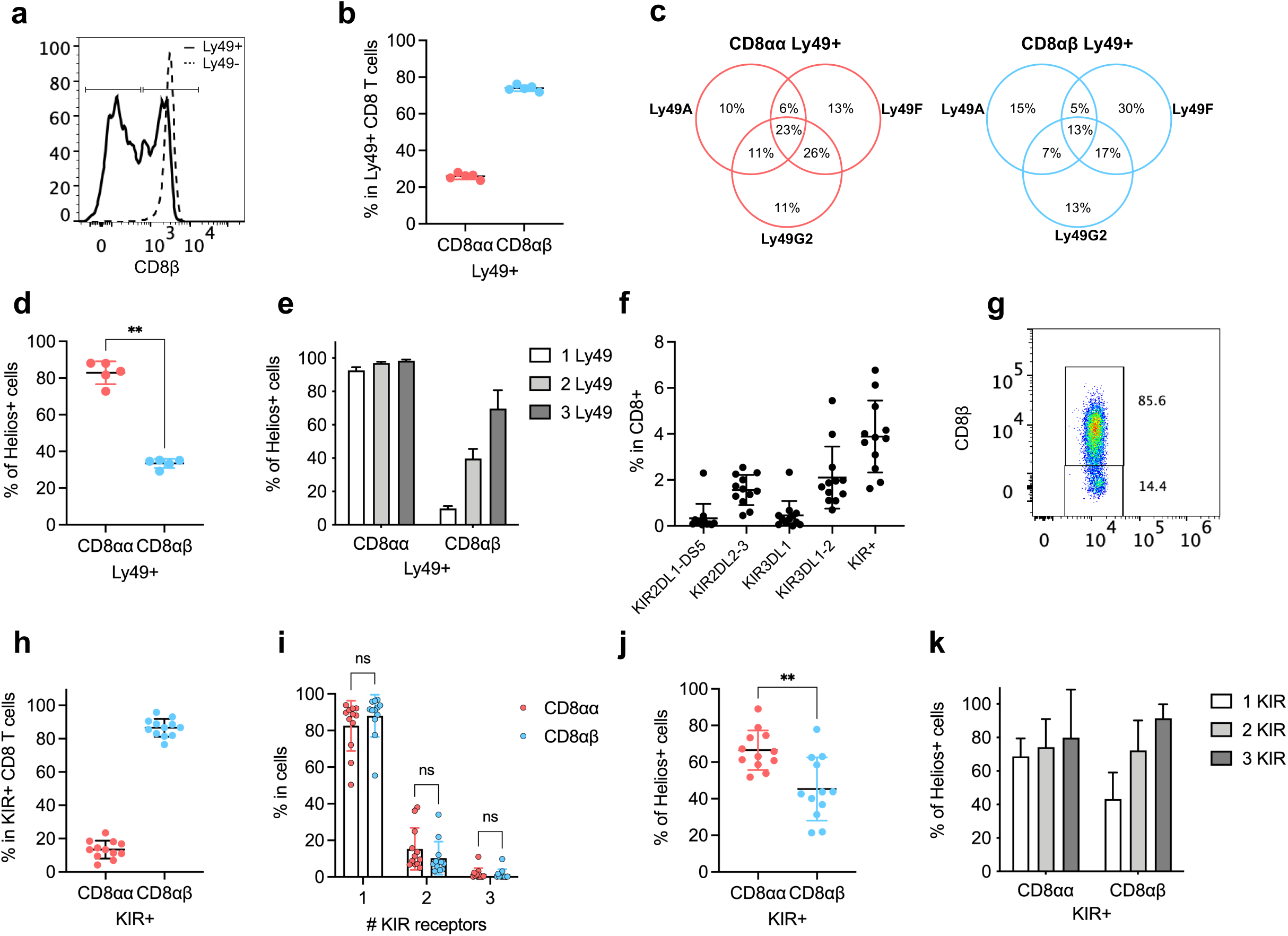
CD8β defines two subsets of murine Ly49+ and human KIR+ CD8 T cells. (**a**) Expression of CD8β by murine CD8 T cells subsets. (**b**) Proportion of CD8⍺⍺-expressing and CD8αβ-expressing cells within Ly49+ CD8 T cells. (**c**) Proportion of each Ly49 receptors combination within Ly49+ subsets and represented as Venn-diagram. (**d**) Helios expression by CD8⍺⍺-Ly49+ and CD8⍺β-Ly49+ cells. (**e**) Helios expression according to the number of Ly49 receptors expressed. (**f**) Expression of inhibitory KIR receptors by CD8 T cells from human PBMCs. (**g**) Expression of CD8β by human KIR+ CD8 T cells. (**h**) Proportion of CD8⍺⍺-expressing and CD8⍺β-expressing cells within KIR+ CD8 T cells. (**i**) Proportion of cells expressing 1, 2 or 3 different KIR receptors within KIR+ subsets. (**j**) Helios expression by CD8⍺⍺-KIR+ and CD8⍺β-KIR+ cells. (**k**) Helios expression according to the number of KIR receptors expressed. Data are represented as mean ± SD (n= 5 mice/group and 12 healthy donors/group). Statistical significance of differences was determined with Mann-Whitney test (ns = non statistical, * p <0.05, ** p <0.01).

KIR+ CD8 T cells are the human counterpart of murine Ly49+ CD8 T cells (Li *et al*., 2022). We thus asked whether CD8αα-expressing and CD8αβ-expressing cells can be found within human KIR+ CD8 T cells. We first analyzed the expression of several inhibitory KIR receptors by CD8 T cells, in the peripheral blood of healthy donors. KIR+ CD8 T cells represent around 4% of total CD8 T cells with KIR3DL1/KIR3DL2 and KIR2DL2/KIR2DL3 being the most expressed (Fig. 1f). There was almost no KIR2DL1/KIR2DS5 expression, hence, it was excluded from the following analysis. Both CD8αα-expressing and CD8αβ-expressing cells were identified within KIR+ CD8 T cells, in proportion equivalent to murine Ly49+ CD8 T cells (Fig. 1g-h). Unlike murine Ly49+ CD8 T cells, CD8αα-KIR+ and CD8αβ-KIR+ cells share the same KIR receptor expression-pattern and express mainly 1 KIR receptor, corresponding to KIR3DL1/KIR3DL2 (Fig. 1i). However, similar to murine Ly49+ CD8 T cells, Helios expression is enriched within CD8αα-KIR+ cells (Fig. 1j) and its expression correlates with the number of KIR receptors expressed within CD8αβ-KIR+ cells only (Fig. 1k). In conclusion, CD8β expression defines two subsets of murine Ly49+ and human KIR+ CD8 T cells which differ in their Ly49/KIR receptor expression pattern and in the expression of the transcription factor Helios.

### In silico lineage tracing of Ly49-expressing CD8 T-cells in the thymus

Multiple lines of evidence point to a thymic origin for Ly49+ CD8 T cells, however their development pathway remains elusive. We analyzed Ly49 receptors expression by thymocytes, after excluding unconventional T cells such as NKT, MAIT or γδ T cells. We observed a small fraction of thymocytes expressing the Ly49A, Ly49F or Ly49G2 receptors (Supplementary Fig. 2a). In contrast to conventional-T-cells-precursor-thymocytes, Ly49+ thymocytes are mainly composed of double negative cells (DN - 45%), CD8 single-positive cells (SP - 35%) and CD4 SP cells (15%) with few double positive (DP) cells (Fig. 2a-c). Moreover, the majority (90%) of Ly49+ DN thymocytes express a TCRβ chain (Fig. 2b-c), compared to less than 10% in total DN (Fig. 2b). A large fraction of those Ly49+ DN TCRβ+ are characterized by the expression of CD5 and to a lesser extent CD69, whose expression is induced following TCR stimulation (Swat *et al*., 1993; Azzam *et al*., 1998), along with the absence of CD24 expression, which is indicative of a mature phenotype (Supplementary Fig. 2b). Similar to total thymocytes, the number of Ly49+ thymocytes increases with age, reaching its maximum around 10 weeks of age (Supplementary Fig. 2c).

**Figure 2:**
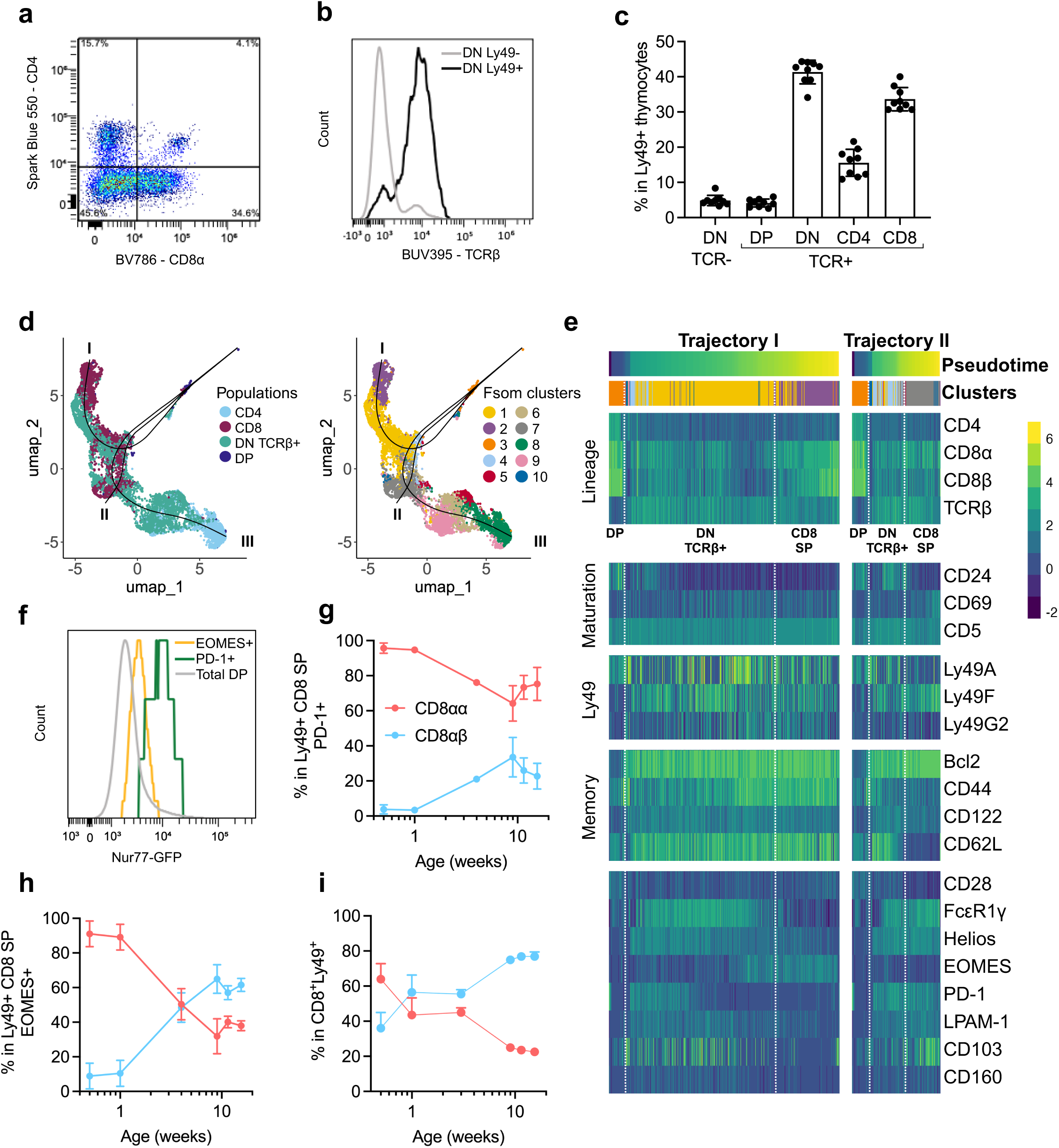
Lineage tracing of Ly49+ thymocytes. (**a**) CD4 and CD8α expression by Ly49+ thymocytes. (**b**) Expression of TCRβ by Ly49+ and Ly49-DN thymocytes. (**c**) Proportion of DN, DP and SP cells among TCR+ and TCR-Ly49+ thymocytes. (**d**) Ly49+ populations or clusters determined by FlowSOM in UMAP space. Slingshot trajectories (black lines) are overlaid on the UMAP plots. (**e**) Heatmap of markers expression along the pseudotime for the trajectories I and II. (**f**) Nur77 (Nr4a1-GFP) expression by EOMES+ and PD-1+ Ly49+ CD8 SP thymocytes as well as total DP. (**g-i**) Evolution of CD8αα-expressing and CD8αβ-expressing cells proportions within PD-1+ Ly49+ CD8 SP thymocytes (**g**), EOMES+ Ly49+ CD8 SP thymocytes (**h**) and splenic Ly49+ CD8 T cells (**i**). Data are represented as mean ± SD (n = 5 mice/group).

This unconventional representation of DN TCRβ+ cells within Ly49+ thymocytes suggest that they could follow an alternative selection process referred to as agonist selection, similar to that of unconventional T cells such a MAIT, NKT or CD81Z1Z-intra-epithelial lymphocytes (IELs). For example, NKT17 and some NKT1 precursors (Hogquist and Georgiev, 2020), as well as some MAIT precursors (Koay *et al*., 2016), lose both CD4 and CD8 expression during their thymic development. Furthermore, CD81Z1Z-IELs are derived from two DN-TCR+ precursors (Klose *et al*., 2014; McDonald *et al*., 2014; Ruscher *et al*., 2017). To test this hypothesis, we performed multiparametric staining of Ly49+ TCR+ thymocytes and performed a pseudotime ordering using Slingshot (Street *et al*., 2018). The markers used were selected for their ability to distinguish the differentiation and maturation stages of thymocytes, and to discriminate memory-phenotype (MP) cells, as well as IELs precursors cells. Following dimensionality reduction of the data, cell clusters were determined using FlowSOM (Fig. 2d). Cluster 3, corresponding to DP cells, was used as the starting cluster for the pseudotime ordering (Fig. 2d). Three trajectories were identified, two leading to CD8 SP cells ending in cluster 2 and 7 and one leading to CD4 SP cells ending in cluster 8 cells (Fig. 2d, Supplementary Fig. 3c). We focused on CD8 SP thymocytes and analyzed the expression of the different markers along the inferred trajectories I and II (Fig. 1e and Supplementary Fig. 3a). Following the DP stage, cells from both trajectories lost CD4 and CD8 expression, and appeared as DN cells. The cells then reacquired CD8α expression, along with CD8β for some cells of the trajectory I. Cells in the cluster 2 and the cluster 7, which were respectively located at the end of the differentiation trajectories I and II, both expressed CD5 and CD69 and low levels of CD24, indicative of a mature phenotype (Fig. 1e and Supplementary Fig. 3 a). However, these two clusters exhibited distinct phenotype. First, cells in cluster 7 primarily expressed the Ly49F receptor, while cells in cluster 2 exhibited a more diverse Ly49 expression pattern, expressing one or more of the three Ly49 receptors. Moreover, cells in cluster 2 had a memory phenotype, characterized by the expression of Bcl2, CD44 and CD122 while cells in cluster 7 had a naive phenotype. Finally, cluster 2 cells were characterized by the expression of Helios and EOMES transcription factors and CD62L homing receptor, while cluster 7 cells were distinguished by the expression of FcεR1γ, Helios, PD-1 and CD103 and LPAM-1 homing receptor.

Thus, our results predicted that Ly49+ thymocytes transition to a DN TCR+ stage after the DP stage and give rise to two subsets of mature Ly49+ CD8 SP thymocytes. Hence, based on EOMES and PD-1 expression, Ly49+ CD8 SP thymocytes can be subdivided in 3 populations that are EOMES+, PD-1+ or EOMES^-^PD-1 (Supplementary Fig. 3b). The EOMES+ subset has a memory phenotype similar to MP-Ly49+ CD8 T cells in the periphery, while the PD-1+ one displays a phenotype reminiscent of the “type A” IELs precursor (Ruscher *et al*., 2017). As previously described (Ruscher *et al*., 2017), these cells expressed high levels of Nur77 (Supplementary Fig. 3c).

We then analyzed the number and proportion of CD8αα-expressing and CD8αβ-expressing cells within EOMES+ and PD-1+ subsets, during post-natal thymic development. At all-time points, the PD-1+ subset comprises almost exclusively CD8αα-expressing cells (Fig. 2g), in line with their IEL precursor potential. In contrast, the relative proportion of CD8αα-expressing and CD8αβ-expressing cells within the EOMES+ subset changes over time. Indeed, during the first week of age, CD8αα-expressing cells represents 90% of the EOMES+ subset (Fig. 2h). Their proportion then declines to reach 30% at 10 weeks of age, while the proportion of CD8αβ-expressing cells increases to reach 70% of the EOMES+ subset (Fig. 2h). These changes in proportions mirrored those of MP-CD8αα-Ly49+ and MP-CD8αβ-Ly49+ T cells within the spleen (Fig. 2i). Although, in the spleen, both MP-CD8αα-Ly49+ and MP-CD8αβ-Ly49+ cell numbers increased with age (Supplementary Fig. 3d). These data are in agreement with our hypothesis that the EOMES+ subset gives rise to the MP-Ly49+ CD8 T cells found in secondary lymphoid organs, whereas the PD-1+ subset differentiates into Ly49+ CD8αα-IELs.

Altogether, our results suggest that following agonist selection, a subset of CD8 SP Ly49+ EOMES+ thymocytes is generated, which could subsequently give rise to both MP-CD81Z1Z-Ly49+ and MP-CD81Zβ-Ly49+ cells in the periphery.

### MP-Ly49+ CD8 T cells require Zeb1 for their development

Since our results suggested similarities in the developmental mechanisms leading to Ly49+ CD8 T cells and iNKT cells, we wanted to further address this issue. A previous report showed that *Cellophane* mutant mice, which bear Zeb1 hypomorphic mutation (Arnold *et al*., 2012), have an altered number of iNKT and MP-Ly49+ CD8 T cells (Zhang *et al*., 2021). Zeb1 attenuates TCR signaling by repressing a signaling module thus helping unconventional T cells, such as NKT cells, to escape negative selection (Zhang *et al*., 2021). We thus wanted to determine the impact of Zeb1 deletion on the development of MP-Ly49+ CD8 T cells in the thymus. Both Ly49+ and Ly49-thymocytes numbers were strongly reduced in Cellophane mice compared to WT (Fig. 3a). Within Ly49+ thymocytes, the proportion of CD8 SP cells was mainly affected in Cellophane mice compared to WT (Fig. 3b). In the periphery, compared to conventional Ly49-CD8 T cells, we observed a strong reduction in both the percentages and numbers of MP-CD8αα-Ly49+ and MP-CD8αβ-Ly49+ cells in Cellophane mice, with MP-CD8αα-Ly49+ cells being the most affected (Fig. 3c-d). Thus, MP-Ly49+ CD8 T cells require Zeb1 for their development. This reinforces our previous hypothesis that these cells are selected in the thymus following agonist selection, similar to iNKT cells.

**Figure 3:**
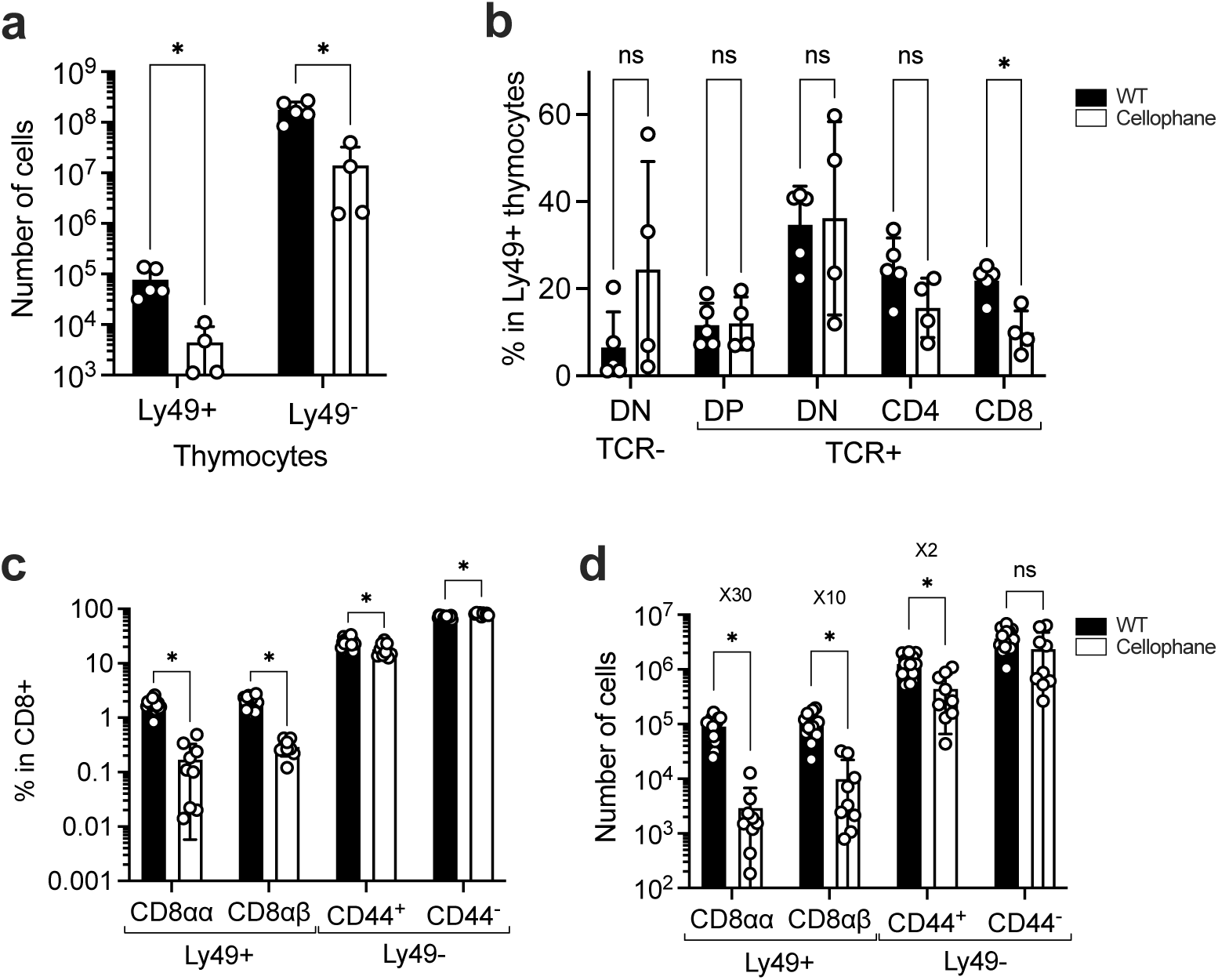
MP-Ly49+ CD8 T cells require Zeb1 for their development. (**a**) Number of Ly49+ and Ly49-thymocytes in WT and Cellophane mice. (**b**) Proportion of thymic subsets within Ly49+ thymocytes of WT or Cellophane mice. (**c-d**) Proportion (**c**) and number (**d**) of CD8 T cells in the spleen of WT and Cellophane mice. Data are represented as mean ± SD (n = 4 to 17 mice/group). Statistical significance of differences was determined with Mann-Whitney test (* p <0.05).

### MP-Ly49+ CD8 T cells participate in the antigen specific antiviral response

We next explored the role of MP-CD81Z1Z-Ly49+ and MP-CD81Zβ-Ly49+ cells in the antiviral response. Following intranasal infection with vaccinia virus (VV), we observed increased numbers of conventional effector CD8 T cells as well as both MP-CD8αα-Ly49+ and MP-CD8αβ-Ly49+ cell subsets in the spleen and the draining lymph nodes (dLN), at the peak of the CD8 T cell response (Fig. 4a). To confirm that MP-Ly49+ CD8 T cells proliferate in response to viral infection, mice received a BrdU injection at 7 days-post-infection (dpi) and the incorporation of BrdU was measured at 8 dpi. As expected, a large fraction of conventional effector CD8 T cells proliferated and incorporated BrdU, while there was little BrdU incorporation in CD8 T cells from naive host (Fig. 4b). Importantly, MP-CD8αα-Ly49+ and MP-CD8αβ-Ly49+ cells subsets incorporated BrdU in both the spleen and the dLN of infected host (Fig. 4b), confirming that their increase in numbers was associated with proliferation.

**Figure 4:**
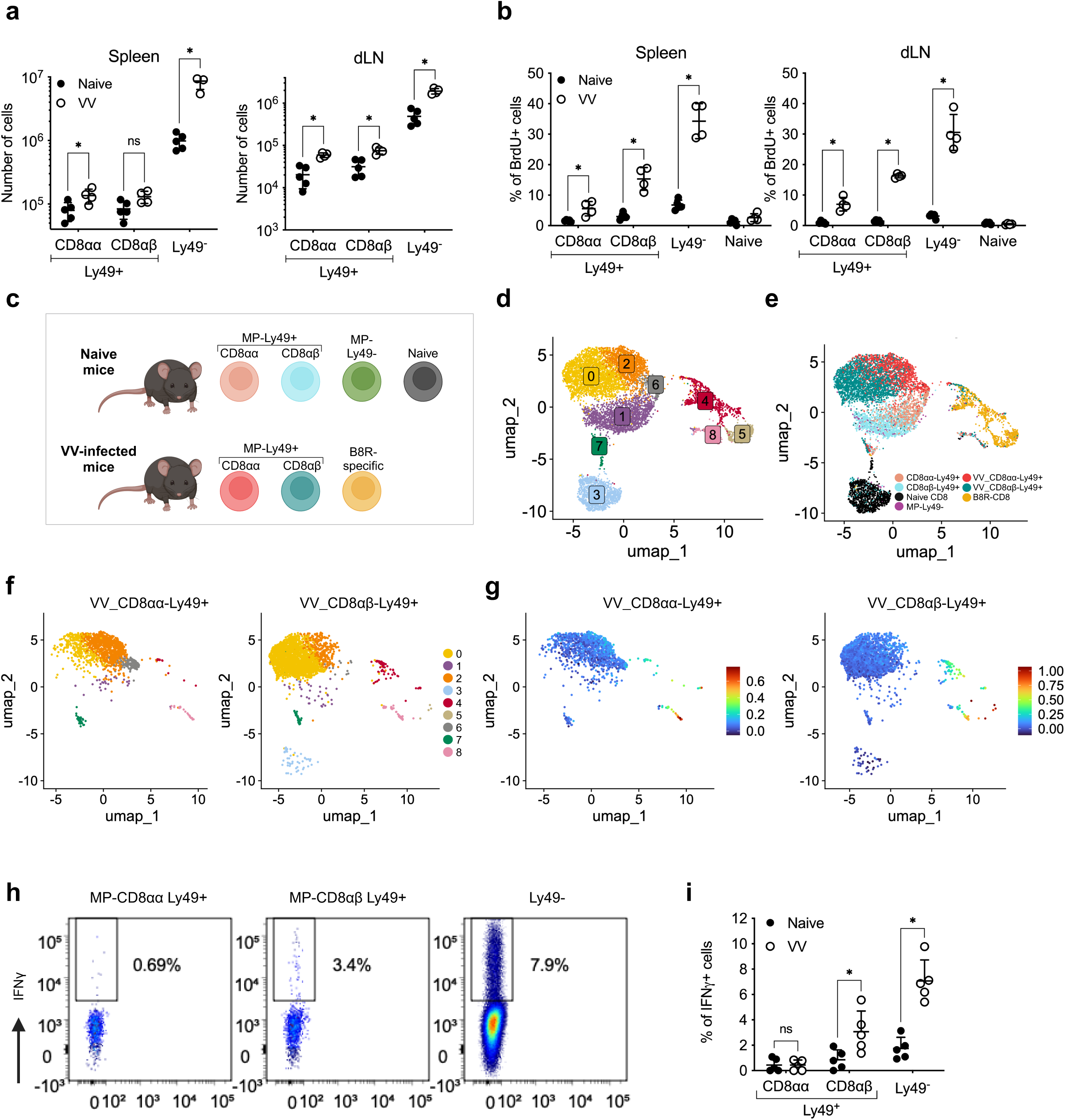
MP-Ly49+ CD8 T cells participate in the antiviral immune response. (**a**) Numbers of CD8 T cells in the spleen and dLN, 8 days after infection with vaccinia virus (VV). (**b**) BrdU incorporation by CD8 T cells from the spleen and dLN of VV-infected mice (8 dpi). (**c**) MPCD8⍺⍺-Ly49+, MP-CD8⍺β-Ly49+, Ly49-CD44+ and naive CD8 T cells were single-cell sorted from the spleen of naive mice. MP-CD8⍺⍺-Ly49+, MP-CD8⍺β-Ly49+ and B8R-specific CD8 T cells were single-cell sorted from the spleen of VV-infected mice (8 dpi). The transcriptome was analysed by scRNA-Seq using 10X technology. Illustration was created with https://BioRender.com. (**d-e**) UMAP plots of the clusters (**d**) and populations (**e**). (**f-g**) MP-CD8αα-Ly49+ and MP-CD8⍺β-Ly49+ cells from VV-infected mice were represented on a UMAP and coloured by clusters (**f**). Enrichment of the effector-gene-expression-signature i.e. the genes up-regulated in B8R-specific cells (cluster 4 and 5) compared to naive cells (cluster 3) (**g**). (**h-i**) Proportion of splenocytes expressing IFNγ following a 4 h *in vitro* stimulation with B8R peptide, 8 dpi with VV. Data are represented as mean ± SD (n = 5 mice/group). Statistical significance of differences was determined with Mann-Whitney test (* p <0.05).

To further characterize the activation status of MP-Ly49+ CD8 T cells during the course of an antiviral response, we performed a single-cell transcriptomic analysis. We sorted both MP-CD8αα-Ly49+ and MP-CD8αβ-Ly49+ cells from naive or VV-infected mice (8 dpi) (Fig. 4c). We also sorted Ly49-CD44+ and naive CD8 T cells from naive mice and VV-epitope-specific CD8 T cells (as revealed by staining with HD^b^-tetramers loaded with the viral peptide B8R, see methods) from VV-infected mice, as controls (Fig. 4c). Clustering analysis partitioned cells into 9 clusters, that can be visualized on a UMAP dimensional reduction (Fig. 4d). Projection of cell types on the UMAP showed that MP-CD8αα-Ly49+ and MP-CD8αβ-Ly49+ cells clustered together, apart from naive and B8R-specific CD8 T cells (Fig. 4e). MP-CD8αα-Ly49+ and MP-CD8αβ-Ly49+ cells from naive mice were located in cluster 1 while MP-CD8αα-Ly49+ and MP-CD8αβ-Ly49+ cells from VV-infected mice were split in two clusters, number 2 and 0 respectively. Naive cells were located in cluster 3 and B8R-specific CD8 T cells were split into clusters 4 and 5. Finally, cluster 6 were enriched in MP-CD8αα-Ly49+ cells from both naive and VV-infected mice, while clusters 7 and 8 contain a mix of MP-Ly49+ CD8 T cell subsets (Fig. 4d-e, Supplementary Fig. 4a).

Although the majority MP-Ly49+ CD8 cells were associated with cluster 2 or 0, some of them and especially MP-CD8αβ-Ly49+ T cells were located in clusters 4 and 5, along with the B8R-specific cells (Fig. 4f, Supplementary Fig. 4b). MP-Ly49+ CD8 T cells in cluster 4 were enriched in genes that are differentially expressed by B8R-specific cells compared to naive CD8 T cells in particular *Gzma*, *Gzmb* or *Lgals3* as well as genes related to proliferation (Fig. 4g, Supplementary Fig. 4c, Supplementary Table 1) suggesting that they respond through their TCR, similarly as B8R-specific T cells.

To confirm the capacity of MP-Ly49+ CD8 T cells to be activated through their TCR, we stimulated splenocytes from VV-infected mice with B8R peptide at 8 dpi. Consistent with the single-cell sequencing results, a significant fraction of MP-CD8αβ-Ly49+ cells from infected mice were able to produce IFNγ in response to B8R stimulation, although this frequency was lower than that of conventional CD8 T cells (Fig. 4h, i). However, the response of MP-CD8αα-Ly49+ from infected or naive control mice were equivalent (Fig. 4h, i), indicating a lack of responsiveness of these cells to TCR triggering via peptide stimulation.

Finally, to determine if MP-Ly49+ CD8 T cells could also respond to a tumoral challenge, B6 mice were inoculated subcutaneously with the EL4 lymphoma cell line expressing NP68 or with VV-NP68 as a control. Mice also received Ly49-CD8 T cells from F5-TCR-transgenic mice, as a tracer for the response to NP68 peptide. We showed that the number of MP-Ly49+ CD8 T cells were increased in the blood of both group of mice, whether challenged by a virus or a tumour (Supplementary Fig. 5a) and the numbers of both MP-CD8αα-Ly49+ and MP-CD8αβ-Ly49+ at 8 dpi were increased in the dLN, but not the spleen of tumor-bearing mice as compared to naive mice (Supplementary Fig. 5b). Finally, we showed that MP-CD8αβ-Ly49+ but not MP-CD8αα-Ly49+ T cells were able to produce IFNγ in response to a stimulation with the NP68 peptide, following both a NP68-viral or NP68-tumoral challenge (Supplementary Fig. 5c).

Overall, our results indicate that in contrast to MP-CD8αα-Ly49+ T cells, MP-CD8αβ-Ly49+ T cells participate in antiviral or antitumoral immune responses, acquire an effector gene-expression signature and are able to respond to TCR-mediated activation via peptide stimulation.

### Viral infection induces a bystander response of MP-Ly49 CD8 T cells

ScRNA-seq analyses revealed that MP-CD8αα-Ly49+ and MP-CD8αβ-Ly49+ T cells had a different gene expression pattern in VV-infected mice than in naive mice. We first determined the gene that are differentially expressed (DE) between MP-CD8αα-Ly49+ and MP-CD8αβ-Ly49+ cells obtained from naive or VV-infected mice. In naive mice, MP-CD8αα-Ly49+ cells were enriched in genes such as *Ikzf2* (Helios), *Fcer1g* (Fcεr1γ), *Itga4* (CD49d), *Klrk1* (NKG2D) and *Cd160* while MP-CD8αβ-Ly49+ cells were enriched in *Il6ra* or *Gzmm* expression (Fig. 5a Supplementary Table 2). These differences were maintained and amplified when cells were isolated from an infected host (Fig. 5b, Supplementary Table 3). We confirmed at the protein level that MP-CD8αα-Ly49+ cells were enriched in Fcεr1γ, CD49d, NKG2D and CD160 expression as compared to MP-CD8αβ-Ly49+ cells (Fig. 5c).

**Figure 5:**
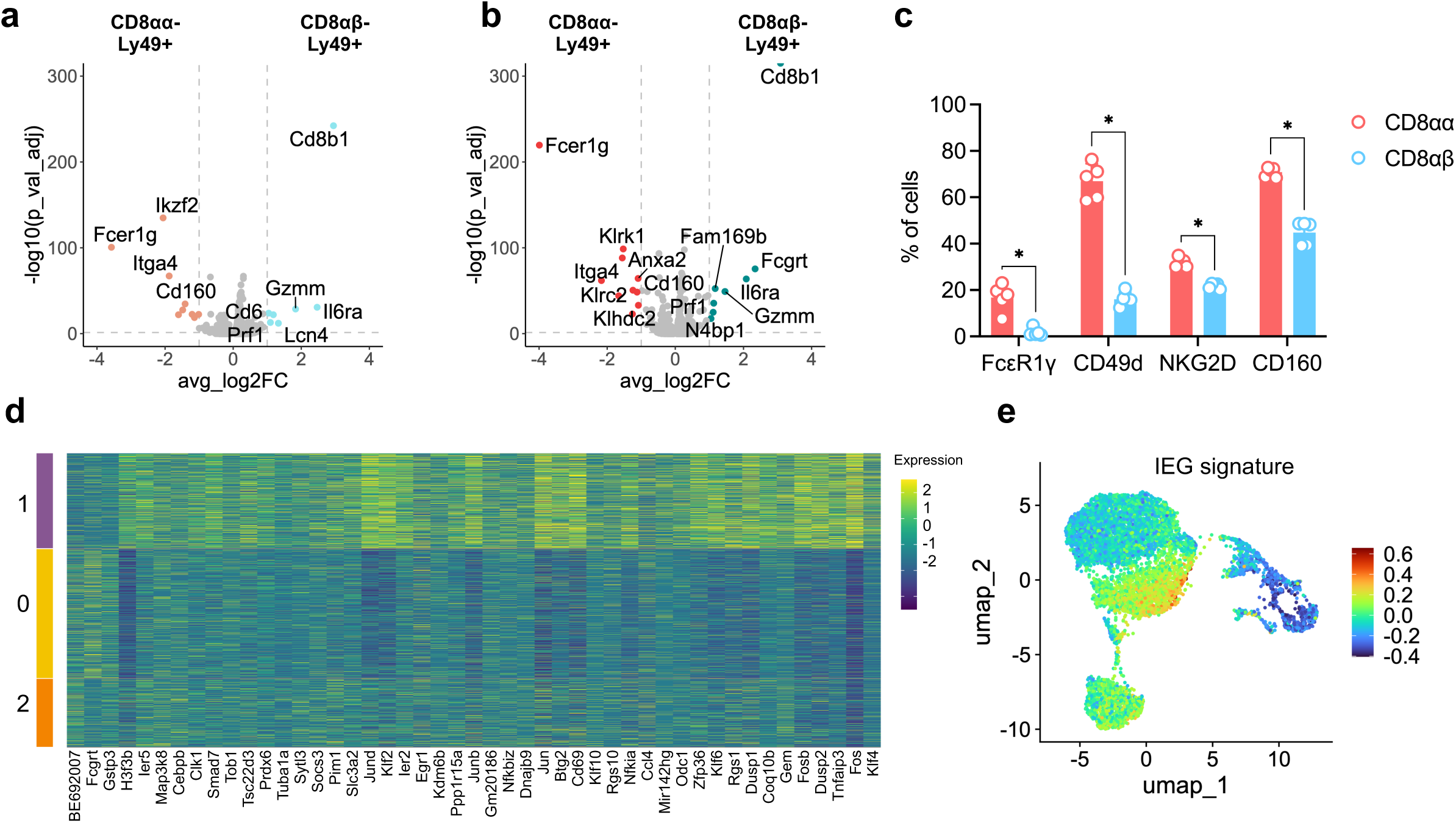
MP-Ly49+ CD8 T cells are enriched in IEG. (**a-b**) Volcano plots of the genes differentially expressed between MP-CD8⍺⍺-Ly49+ and MPCD8⍺ β-Ly49+ cells from naive (**a**) and VV-infected (**b**) mice. (**c**) Expression of FcεR1γ, CD49d, NKG2D, CD160 and IL6 receptor by MP-CD8⍺⍺-Ly49+ and MP-CD8⍺β-Ly49+ cells in the spleen of naive mice, measured by flow cytometry. (**d**) Heatmap of the genes differentially expressed between cells from cluster 1 versus cells from clusters 0 and 2. (**e**) UMAP plot coloured according to the IEG signature. Data are represented as mean ± SD (n = 5 mice/group). Statistical significance of differences was determined with Mann-Whitney test (* p <0.05).

We then compared the DE genes between MP-Ly49+ CD8 T cells from cluster 1 (Naive) and clusters 0 and 2 (VV-infected). Following viral infection, 3 genes (*Fcgrt*, *BE692007, GSTP3*) were upregulated in MP-Ly49+ CD8 T cells from infected host, and 44 genes were downregulated (Fig. 5d). Most of those genes, such as *Egr1*, *Fos* or *Jun,* belong to the family of immediate early genes (IEG). Indeed, MP-Ly49+ CD8 T cells were enriched in the IEG signature as compared to naive mice, and this signature is downregulated in infected host (Fig. 5e). A similar downregulation is observed in B8R-specific CD8 T cells compared to MP-Ly49+ CD8 T cells.

Altogether, our results indicates that during a course of a viral infection, MP-Ly49+ CD8 T cells modify their gene expression profile without acquiring genes related to TCR stimulation. This suggest that these cells respond to cytokine-driven bystander stimuli.

### MP-Ly49+ CD8 T cells are highly sensitive to cytokine stimulation

To identify the cytokines that could drive the bystander activation of MP-Ly49+ CD8 T cells, we first analyzed the expression of Stat genes in our scRNA-Seq data. MP-Ly49+ CD8 T cells were enriched in *Stat3* and more importantly *Stat4* expression compared to naive or B8R-specific CD8 T cells (Fig. 6a). We then analyzed the expression of genes coding for cytokine receptors. At the gene level, MP-Ly49+ CD8 T cells were particularly enriched in *Ifngr1*, *Il2rb*, *Il7r*, *Il17ra* and *Il18r1* expression (Fig. 6b). Furthermore, they expressed higher level of *IL4ra* and *Il10rb* than both naive and B8R-specific CD8 T cells (Fig. 6b) and MP-CD8αβ-Ly49+ cells were enriched in *Il6ra* expression (Fig. 5c-d, Fig. 6b). These differences were confirmed at the protein level with both MP-CD8αα-Ly49+ and MP-CD8αβ-Ly49+ cells expressing higher levels of CD122 (IL-15Rβ), CD124 (IL-4Rα), CD212 (IL-12Rβ1) and CD218 (IL-18Rα) compared to naive or MP-Ly49-CD8 T cells (Fig. 6c). All subsets maintained equivalent levels of CD127 (IL-7R) while MP-CD8αβ-Ly49+ cells displayed a slight increase in the expression level of CD126 (IL-6Rα) compared to other subsets (Fig. 6c).

**Figure 6:**
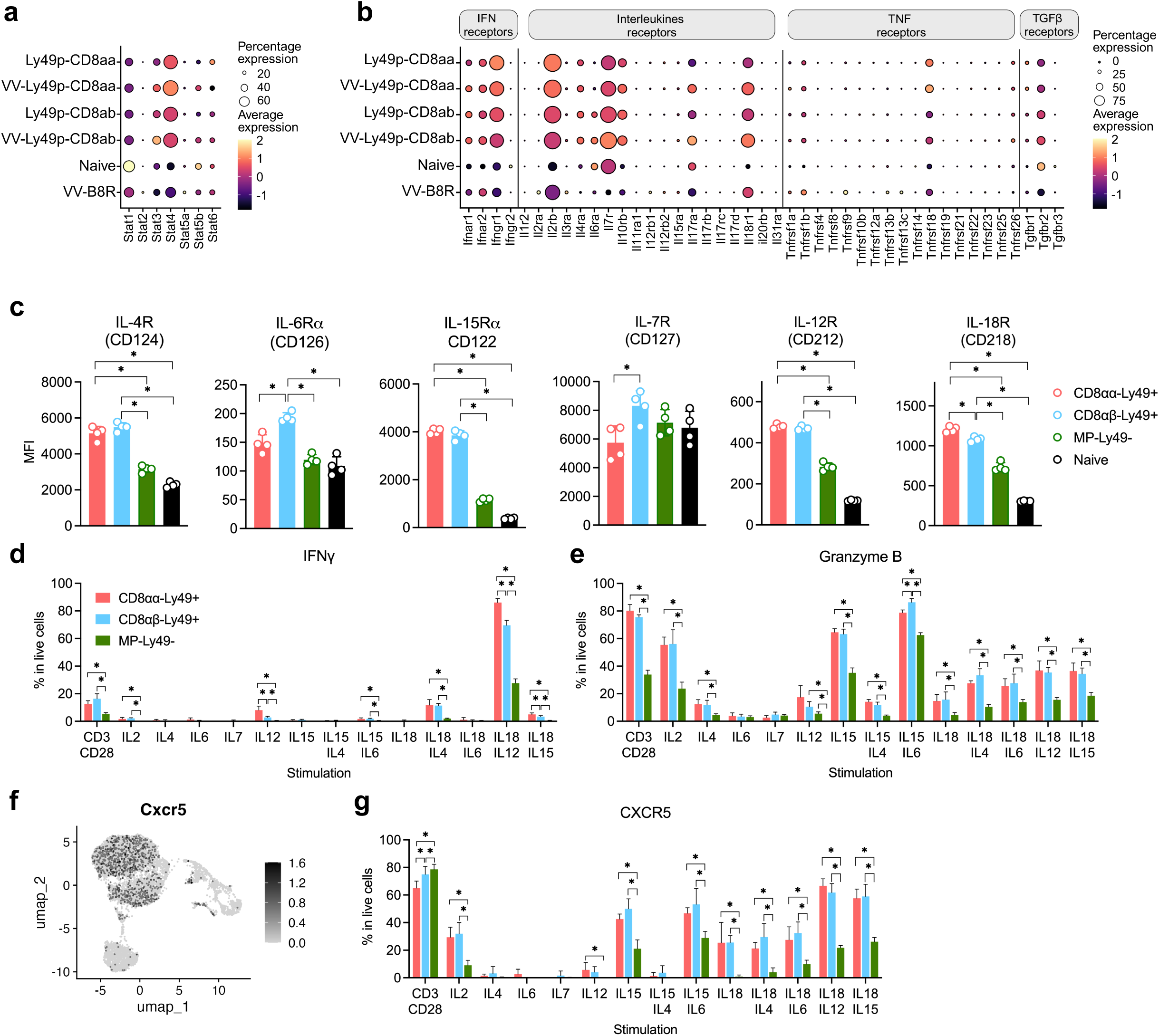
MP-Ly49+ CD8 T cells are highly sensitive to cytokine stimulation. (**a**) DotPlot of Stat genes expression. (**b**) DotPlot of the expression of the genes coding for cytokines receptors. (**c**) Expression of IL receptors by CD8 populations, determined by flow cytometry and expressed in MFI. (**d-e**) Expression of IFNγ (**d**) and granzyme B (**e**) by MP-CD8 T cells, 24h after in vitro stimulation, and normalized against unstimulated control. (**f**) UMAP plot of the expression of the Cxcr5 gene. (**g**) Expression of CXCR5 by MP-CD8 T cells, 24h after in vitro stimulation, and normalized against unstimulated control. Data are represented as mean ± SD (n = 5 mice/group).

In conventional CD8 T cells, bystander activation can lead to the expression of IFNγ or granzyme B. We thus addressed the impact of the cytokines binding the receptors expressed by MP-Ly49+ CD8 T on the expression of both proteins. TCR/CD28 stimulation was used as a control. Around 15% of MP-Ly49+ CD8 T cells were able to produce IFNγ in response to TCR/CD28 stimulation (Fig. 6d). There was also a weak IFNγ production by MP-Ly49+ CD8 T cells in response to IL-2, IL-12, IL-15 combined with IL-6, and IL-18 combined with IL-4 or IL-15 (Fig. 6d). More importantly, there was a massive IFNγ production by MP-Ly49+ CD8 T cells in response to IL-18 combined with IL-12, compared to MP-Ly49-CD8 T cells (Fig. 6d). This IL-12/IL-18-driven IFNγ response was higher for MP-CD8αα-Ly49+ cells, in agreement with their higher level of expression of CD218 (Fig. 6c-d).

Granzyme B was also induced by TCR/CD28 stimulation, as well as by IL-2 or IL-15 alone or in combination with IL-6. Interestingly, the addition of IL-4 to IL-15 strongly inhibited the expression of granzyme B (Fig. 6e).

To exert their regulatory function on Tfh, MP-Ly49+ CD8 T cells have to migrate to the germinal centers in the LN, which requires the expression of the chemokine receptor CXCR5 (Kim *et al*., 2011). In the scRNAseq analysis, both MP-CD8αα-Ly49+ and MP-CD8αβ-Ly49+ cells were enriched in *Cxcr5* gene expression, compared to naive or B8R-specific CD8 T cells (Fig. 6f). We measured the impact of cytokine stimulation on CXCR5 expression. Following TCR/CD28 stimulation, we observed a massive upregulation of CXCR5 expression by MP-Ly49+ as well as MP-Ly49-CD8 T cells (Fig. 6g). Moreover, CXCR5 was upregulated on MP-Ly49+ CD8 T cells in response to IL-2, IL-12, IL-15 alone or combined with IL-6, and IL-18 alone or in combination with various cytokines (Fig. 6g). As observed for granzyme B expression (Fig. 6e), IL-4 antagonized the effect of IL-15 on CXCR5 expression (Fig. 6g), indicating an inhibitory role for this cytokine.

In conclusion, bystander activation of MP-Ly49+ CD8 T cells by a number of cytokines, induces the expression of effector molecules such as IFNγ or GZMB and facilitates the expression of CXCR5, that is key to access germinal centers.

## Discussion

Our study identifies two subsets of murine Ly49+ and human KIR+ MP CD8 T cells, based on the expression of the CD8β chain. In both subsets, CD8αα-expressing cells are enriched in Helios expression. Furthermore, in mice, MP-CD8αα-Ly49+ cells can express multiple Ly49 receptors simultaneously, while MP-CD8αβ-Ly49+ cells primarily express the Ly49F receptor. In human, both subsets express mainly a single KIR receptor. Similar to MP CD8 T cells, MP-CD8αβ-Ly49+ express low levels of CD49d (Haluszczak *et al*., 2009) and NKG2D (Ventre *et al*., 2012; Grau *et al*., 2018), unlike MP-CD8αα-Ly49+ cells. Furthermore, MP-CD8αα-Ly49+ cells are enriched in *Fcer1g* expression while MP-CD8αβ-Ly49+ cells are enriched in *Il6ra* and *Gzmm* expression.

Following activation, CD8 T cells acquire the expression of NK cell receptors, such as NKG2D, whose expression is maintained on memory cells (Grau *et al*., 2018). Unlike these receptors, Ly49 proteins expression is not induced on naive CD8 T cells, whether *in vitro* (Anfossi *et al*., 2004) or *in vivo*, in different stimulation contexts such as cytokine stimulation, lymphopenia (McMahon *et al*., 2002) or viral infection (Peacock *et al*., 2000; Peacock and Welsh, 2004). This suggests that these receptors are acquired by CD8 T cells during thymic differentiation. To investigate the thymic origin of MP-Ly49+ CD8 T cells, we employed a trajectory inference tool based on surface protein and transcription factors expression. Our analysis predicted that following the DP stage, Ly49+ thymocytes loose the expression of both CD4 and CD8 coreceptors and emerge as DN cells expressing a TCR. These DN TCR+ cells then diverge into three distinct branches. One branch differentiates into Ly49+ CD4 T cells, predominantly expressing the Ly49A receptor. These cells potentially correspond to α-GalCer:CD1d unrestricted NKT (Farr *et al*., 2014). The remaining two branches give rise to CD8 SP subsets, one of which expresses EOMES and the other expresses PD-1.

This differentiation path is reminiscent of the one described for TCRαβ IELs. Indeed, following negative selection, IEL precursors lose the expression of CD4 and CD8 coreceptors and differentiate into two DN TCR+ precursors, termed “type A” and “type B” (Ruscher *et al*., 2017). Type A precursors exhibit a naive profile and express PD-1 and LPAM-1 while type B precursor are characterized by the expression of Tbet, NK1.1 and CD103 (Ruscher *et al*., 2017). Consequently, the PD-1+ Ly49+ SP thymocyte subset identified in our study, which predominantly include CD8αα-expressing cells, could correspond to “type A” IEL precursors. The “type B” precursors, characterized by NK1.1 expression, would have been excluded from our analysis.

The EOMES+ thymocyte subset mirrors the phenotype of MP-Ly49+ CD8 T cells found in the periphery. This subset exhibits a memory phenotype, characterized by high levels of CD44, CD122, Bcl2, and CD62L and express multiple Ly49 receptors as well as the Helios transcription factor. Notably, the EOMES+ thymocyte subset includes both CD8αα-expressing and CD8αβ-expressing cells, in proportions similar to those of peripheral MP-Ly49+ CD8 T cells, regardless of the mouse’s age. Thus, our findings suggest that both subsets of MP-Ly49+ CD8 T cells may derive from the EOMES+ Ly49+ CD8SP thymocyte subset. This is consistent with a recent study showing that the development of MP-Ly49+ CD8 T cells is under the control of EOMES (Mishra *et al*., 2021).

The decision between the EOMES+ or PD-1+ lineage could be instructed by the TCR. Indeed, we showed that the PD-1+ subset exhibits higher levels of Nur77 compared to the EOMES+ subset, suggesting that stronger TCR signaling favors differentiation into the PD-1+ subset. Additionally, TCR avidity could influence whether cells in the EOMES+ subset express a CD81Z1Z or a CD81Zβ coreceptor. Indeed, it was showed that high doses of agonist peptides select CD8αα-expressing cells while low doses select CD8αβ-expressing cells (Yamagata, Mathis and Benoist, 2004).

Our results, consistent with the existing literature, support the notion of an alternative thymic selection process for MP-Ly49+ CD8 T cells, known as agonist selection. Evidence supporting this includes the observation that transgenic mice bearing classical MHC-I-restricted TCRs, such as female H-Y (Coles *et al*., 2000; Badr *et al*., 2023), OT-1 (Kim *et al*., 2024) or F5 (data not shown), lack MP-Ly49+ CD8 T cells despite being capable of generating MP-CD8 T cells. In contrast, transgenic mice with a FL9 self-peptide-specific TCR (Kim *et al*., 2024), which is Qa-1-restricted, or male HY (Badr *et al*., 2023), can generate MP-Ly49+ CD8 T cells. Additionally, MP-Ly49+ CD8 T cells express the Helios transcription factor, which is indicative of autoreactivity (Daley, Hu and Goodnow, 2013). Thus, similarly to CD4 Treg (Savage, Klawon and Miller, 2020), MP-CD8 Ly49+ derive from thymic precursor with high self-pMHC ligands affinity. This is consistent with their regulatory properties.

Our data reveal that both MP-CD8αα-Ly49+ and MP-CD8αβ-Ly49+ cells are highly dependent on the transcription factor Zeb1 for their development. Similar to what has been described for NKT thymic development (Zhang *et al*., 2021), Zeb1 may enable Ly49+ thymocytes to escape negative selection by modulating the TCR signals driven by autoreactive TCRs. Ly49 receptors may also play a crucial role in the repression of TCR signaling. Indeed, in mice transgenic for the human KIR2DL3 inhibitory receptor and its cognate HLA-Cw3 class I ligand (KIR-HLA Tg), there is an increased proportion of MP CD8 T cells compared to control mice, and notably MP-Ly49+ CD8 T cells (Ugolini *et al*., 2001). This development depends on KIR2DL3 receptor engagement with its ligand, as it is abrogated in the absence of the HLA-Cw3 ligand (Ugolini *et al*., 2001). Furthermore, in KIR-HLA Tg mice expressing the LCMV-specific P14 TCR, MP-Ly49+ CD8 T cells development is compromised, suggesting that KIR receptor signaling alone is insufficient for their development. Our results showed that DP Ly49+ cells were particularly enriched in Ly49A expression. Interestingly, in Ly49A-transgenic mice expressing its main ligand H2-D^d^, the development of autoreactive T cells is exacerbated (Fahlén *et al*., 2000; Pauza *et al*., 2000). Thus, the inhibitory signals provided by Ly49 receptors and especially Ly49A, could play a critical role in modulating TCR signaling and enabling the development of MP-Ly49+ CD8 T cells through agonist selection. In the periphery, the engagement of inhibitory Ly49 receptors, such as Ly49G2, suppresses the cytotoxic activity of Ly49+ MP-CD8 cells, further confirming the capacity of these receptors to downregulate TCR signaling in CD8 T cells (Peacock *et al*., 2000; Peacock and Welsh, 2004).

Despite their few phenotypic differences, MP-CD8αα-Ly49+ and-MP-CD8αβ-Ly49+ cells shared an enrichment in the expression of genes belonging to the IEG family, such as *Jun* or *Fos*. This may reflect the high TCR signals experienced by these cells during their agonist selection (Chopp *et al*., 2020) or could result from enhanced stimulation in the periphery through tonic TCR signaling (Myers, Zikherman and Roose, 2017; Eggert and Au-Yeung, 2021). Alternatively, cytokines such as IL-15 also promote the expression of IEG such as *Fos* or *Jun* through the MAPK signaling pathway (Ma, Caligiuri and Yu, 2022).

In this study, we wanted to address the stimuli that initiate MP-Ly49+ CD8 T cells response in a context of viral infection. We showed that both MP-CD8αα-Ly49+ and MP-CD8αβ-Ly49+ cells proliferate in the spleen and dLN in response to VV infection, corroborating previous findings (Peacock *et al*., 2002; Shytikov *et al*., 2021; Li *et al*., 2022). Our study suggests that the majority of MP-Ly49+ CD8 T cells respond to bystander innate stimuli rather than TCR-mediated activation, as all cells downregulate a certain number of genes, including IEG, in response to viral infection, without acquiring an effector program. Moreover, we showed that a subset of MP-CD8αβ-Ly49+ cells can mediate a classical-MHC-I restricted viral peptide-specific response through TCR signaling, while the MP-CD8αα-Ly49+ did not. This could indicate that MP-CD8αα-Ly49+ recognize different peptides and/or restriction elements.

To identify the factors initiating the MP-Ly49+ CD8 T cell bystander response, we stimulated these cells with various cytokines. We showed that MP-CD8αβ-Ly49+ and especially MP-CD8αα-Ly49+ exhibit heightened sensitivity to *in vitro* IL-12/IL-18 stimulation compared to Ly49-cells, similar to their human KIR+ CD8 T cells counterpart (Jacomet *et al*., 2015). Supporting these results, we found that these cells are enriched in *Stat4* expression (Nakahira *et al*., 2002) and express higher levels of both CD212 and CD218 receptors compared to naive and MP-Ly49-CD8 T cells. MP-Ly49+ CD8 T cells were shown to express CXCR5, facilitating their migration to LN-B cell follicles (Kim *et al*., 2011). Our study indicates that both MP-Ly49+ and MP-Ly49-CD8 T cells upregulate CXCR5 in response to TCR stimulation. More importantly, MP-Ly49+ CD8 T cells upregulate CXCR5 in response to cytokines such as IL-2, IL-15, or IL-18, a response not observed in MP-Ly49-cells. The combination of IL-12 and IL-18 was the most potent stimulus for CXCR5 upregulation. Altogether, these results suggest that during a viral infection, environmental inflammatory cytokines could promote CXCR5 and granzyme B expression in MP-Ly49+ CD8 T cells, enabling their effector function and migration to dLN and germinal centers.

In conclusion, we showed that MP-Ly49+ CD8 T cells is a heterogeneous population comprising both CD8αα-expressing and CD8αβ-expressing cells, that are conserved between species. While these cells exhibit high self-reactivity due to their thymic agonist selection, some of them and especially the CD8αβ-expressing one, may respond to a broad range of antigens, including exogenous antigens. Moreover, the MP-Ly49+ CD8 T cells display heightened sensitivity to bystander stimulation, that could promote their regulatory activity during an immune response. The role in the control of CD4 Tfh cells of the TCR-activated cells compared to the bystander-activated one, remains to be determined.

## Materiel and methods

### Mice

C57Bl/6J mice were purchased from Charles River Laboratories (L’Arbresle, France) and housed under SPF conditions in our animal facility (AniRA-PBES, Lyon, France). F5 TCR-transgenic mice on a B6 background (B6/J-Tg(CD2-TcraF5,CD2-TcrbF5)1Kio/Jmar)(Jubin *et al*., 2012) were crossed with B6-Ptprc^em(K302E)Jmar^/J)(Laubreton *et al*., 2023) to obtain F5 x CD45.1 mice (B6-^Ptprcem(K302E)Jmar^/J-Tg(CD2-TcraF5,CD2-TcrbF5)1Kio/Jmar). Cellophane (Arnold *et al*., 2012) and Nr4a1-GFP (Moran *et al*., 2011) mice were bred and housed under SPF conditions in our animal facility (AniRA-PBES, Lyon, France). Mice aged between 2-days to 20 weeks were used. All experiments were approved by our local ethics committee (CECCAPP, Lyon, France) and the relevant governmental agencies.

### Human samples

Blood samples were collected from healthy donors and provided by the EFS. PBMCs were isolated from whole blood after Ficoll gradient centrifugation (Eurobio Scientific) and cryopreserved.

### Mouse immunization

The recombinant vaccinia virus (VV) expressing the NP68 epitope (VV-NP68) was engineered from the Western Reserve strain by Dr. D.Y.-L. Teoh, in Prof. Sir Andrew McMichael’s laboratory at the Medical Research Council (Human Immunology Unit, Institute of Molecular Medicine, Oxford, U.K.). EL4 lymphoma cell line expressing the NP68 epitope (EL4-NP68) was provided by Dr. T.N.M. Schumacher. For immunization, mice anesthetized by intraperitoneal (ip) injection of ketamine/xylazine (50 and 10 mg/kg respectively) received intranasal (in) instillation of 2.10^5^ pfu of VV-NP68 under 20 μL. For tumoral challenge, mice received a subcutaneous injection with EL4-NP68 cells (2.,5.10^6^ under 200 μL). In some experiments, mice were intravenously transferred with naive F5 x CD45.1 cells (2.10^5^ under 200 μL), one day prior viral or tumoral challenge. To measure cell proliferation, mice received one ip injection of BrdU (5-Bromo-2’-deoxyuridine, Sigma-Aldrich) at 7 days post-infection (dpi) to label proliferating cells.

### Sample collection and flow cytometry analysis

Mice were sacrificed by cervical dislocation. Spleen, draining lymph nodes (cervical and inguinal) and thymus were harvested, mechanically disrupted, and filtered through a sterile 100-mm nylon mesh filter (BD Biosciences). Single cell suspensions were first incubated with a viability Dye (ThermoFischer Scientific or TFS) for 20 min at 4 °C. To reduce non-specific antibody binding, Fc receptors were blocked with the antibody (Ab) 2.4G2, for 10 min at 4 °C. Surface staining was performed with an appropriate mixture of Ab diluted in staining buffer (PBS supplemented with 1% fetal calf serum (BioWest) and 0.09% NaN3 (Sigma-Aldrich)) for 30 min at 4 °C. For intracellular staining, cells were fixed and permeabilized according to the manufacturer’s instructions using either the CytoFix/CytoPerm buffer (BD Biosciences) for cytokines staining or the Fixation/Permeabilization buffer from Foxp3 Transcription Factor Staining Buffer Kit (TFS) for transcription factors staining or the BrdU Staining Kit (TFS). Fixed cells were then stained with an appropriate mixture of intracellular Ab for 30 min at 4 °C. Analysis were performed either on a BD LSRFortessa cell analyzer (BD Biosciences) and further analyzed using FlowJoTM v10 software (BD Biosciences) or on Cytek® Aurora (Cytek) and analyzed using OMIQ software (DotMATICS). Abs used are listed in the Supplementary Table 4.

### In vitro stimulation

Murine primary cells were cultured in complete DMEM consisting of DMEM medium with 4.5 μg/mL of glucose and GlutaMAX™ supplement (TFS), supplemented with 6% Fetal Calf Serum (BioWest), MEM Non-Essential Amino Acids Solution, 50 μg/mL gentamicin, 10 mM HEPES buffer and 50 μM 2-Mercaptoethanol (50 μM) (all from TFS), and maintained at 37 °C in 7% CO2 incubator.

Spleen single-cell suspensions were stimulated with B8R peptide at 10 nM (TSYKFESV, Proteogenix) for 4 hours in the presence of GolgiStop™ (BD Biosciences), according to manufacturer’s instructions and the expression of IFNγ was measured by flow cytometry. CD8 T cells were enriched from spleen single-cell suspensions using CD8α+ T Cell Isolation Kit and autoMACS® Pro Separator according to the manufacturer’s intructions (Milteniy Biotec). Purity was ≥ 90%. Enriched CD8 T cells (2.10^5^) were incubated for 24 hours at 37°C with Dynabeads™ Mouse T-Activator CD3/CD28 (ratio 1:1, Gibco), murine rIL-2 supernatant (5% corresponding to a final concentration of 11.5 ng/mL), IL-4 (20 ng/mL), IL-6 (50 ng/mL), IL-7 (10 ng/mL), IL-12 (10 ng/mL) or IL-15 (20 ng/mL) from Miltenyi Biotech, or IL-18 (10 ng/mL) from R&D systems. GolgiStop™ was added for the last 4 hours of culture, to allow for cytokine production measurement by intracellular staining. Flow-Count Fluorospheres (Beckman Coulter) were added prior staining, to determine absolute cell numbers per well.

### Pseudotime inference

Doublets of thymic cells were excluded by plotting the width and height of the forward and side scatter. Lymphocytes were then gated based on forward and side scatter and dead cells were excluded according to viability staining. NKT, MAIT and γδ T cells were excluded from the analysis using PB-S57/CD1d tetramer, MR1-5-OP-RU tetramer and anti-γδ TCR and anti-NK1.1 antibody labelling. Ly49+ thymocytes were then selected according to their expression of Ly49A, Ly49F or Ly49G2 receptors. Ly49+ thymocytes data were exported as csv files and imported into R studio (v2024.04.0+735). Clustering was performed using FlowSOM(Van Gassen *et al*., 2015) and UMAP dimensionality reduction of the data was applied. Pseudotemporal ordering of cells was performed in Slingshot(Street *et al*., 2018) using FlowSOM clusters and the UMAP coordinates. The cluster corresponding to DP cells was used as a starting point. No ending clusters were defined. Pseudotime-ordered heatmaps were generated using plotHeatmap ().

### Cell sorting and scRNAseq

CD8 T cells were enriched from spleen single-cell suspensions using CD8α+ T Cell Isolation Kit and autoMACS® Pro Separator according to the manufacturer’s intructions (Milteniy Biotech). CD8 T cell subsets were then FACS-sorted from the CD8 enriched-fraction of naive mice (naive, Ly49-CD44+, CD8αα-Ly49+ and CD8αβ-Ly49+ subsets) or VV-infected mice (VV_CD8αα-Ly49+, VV_CD8αβ-Ly49+ and VV_B8R subsets) on an ARIA II (BD Biosciences), according to their expression of CD8α, CD8β, CD44, Ly49A, Ly49F or Ly49G2 receptors or B8R-dextramer (Immudex) labelling. Sorted cells were centrifuged and barcoded with hashtag oligonucleotides (HTO) antibodies specific for each subset (TotalSeq-B – Biolegend, listed in the supplementary Table 5) for 30 min on ice. Barcoded-subsets were then pooled, centrifugated and 50,000 cells were loaded onto the Chromium 39 chip. After reverse transcription, barcoded cDNAs were amplified by PCR and size-selected to separate cell-cDNAs and HTO-cDNAs, according to the the manufacturer’s instructions. Briefly, the cell-cDNA library was fragmented, end-repaired and sample indexes were added by PCR. Sample indexes were added by PCR for the HTO-cDNA library. The purified libraries were quantified using the Library Quantification Sample Kit Kapa (Illumina). mRNA and HTO libraries were pooled in a 4:1 ratio and paired-end sequenced (2 x 150 pb, 375M reads – Macrogen) on a HiSeq X platform (Illumina), with 1% of PhiX.

### scRNA-seq analyses of murine Ly49+ CD8 T cells

Quality control was performed using FastQC. The TSO, poly-A sequences and low-quality bases were trimmed using cutadapt (Kechin et al., 2017). Transcript expression quantification was performed using the alevin-fry pipeline (He et al., 2022) and version M31 of GENCODE mouse genome and annotations. The gene/count matrix was generated with fishpond (Zhu et al., 2019). Cells with nFeature_RNA > 6500, nCount_RNA > 80 000 and percent.mt > 8%were filtered-out. The HTO library was demultiplexed using the Seurat pipeline (v5.1.0) (Hao et al., 2021) and the RNA counts were normalized using the sctransform function (v2). Data were then scaled with the ScaleData() function followed by principal component analysis with the RunPCA() function on variable features. The ElbowPlot() function was used to determine the optimal number of dimensions (11 CP). Clustree (Zappia and Oshlack, 2018) was used to select the clustering resolution (0.4). Cells clusters were identified with the FindCluster function. Populations and clusters were plotted on UMAP using the RunUMAP() function. Differentially expressed genes were identified using the FindMarkers() function, with the following parameters for cluster 1 versus clusters 0 and 2 (logfc.threshold = 0.75, min.pct = 0.25), cluster 3 versus clusters 4 and 5 (logfc.threshold = 2, min.pct = 0.5) and CD8αα-Ly49+ versus CD8αβ-Ly49+ (logfc.threshold = 1, min.pct = 0.25). Heatmaps were built using the DoHeatmap() function. IEG gene signature (Tullai *et al*., 2007) was displayed using the AddModuleScore() function.

### Statistical analysis

Data were expressed as mean ± SEM. Mann-Whitney test (comparison of two groups) was used to compare unpaired values (GraphPad Prism software). Significance is represented: * p < 0.05; ** p < 0.01; *** p < 0.001; and **** p < 0.0001.

## Acknowledgment

We acknowledge the contribution of SFR BioSciences (UAR3444/CNRS, US8/Inserm, École Normale Supérieure de Lyon, Université de Lyon) and of the CELPHEDIA infrastructure (http://www.celphedia.eu/), especially the center AniRA in Lyon (AniRA-Cytométrie and AniRA-PBES facilities). We thank the NIH Tetramer Core Facility (contract number 75N93020D00005) for providing 5-OP-RU_MR1 and CD1d_PBS-S7 tetramers. We thank Antoine Marçais and Olivier Thaunat for critical reading of the manuscript. This work was supported by INSERM, CNRS, Université de Lyon and ENS Lyon. Margaux Prieux has a région Auvergne-Rhône-Alpes PhD fellowship.

## Data availability

The sequencing data generated in this study are available at GEO NCBI under the accession number GSE282481. The flow cytometry data used to determine the thymic differentiation of MP-Ly49+ CD8 T cells generated in this study are available at https://doi.org/10.6084/m9.figshare.27861636.v1.

## Author contribution

D.L. and J.M. conceived the study and designed the experiments. D.L., M.G., M.P. and M.T. performed the experiments. D.L., M.P. and V.M. conducted the bioinformatic analysis. T.W. provide Cellophane mice. D.L. and J.M. wrote the paper. M.G., V.M., M.P., M.T., M.V, and T.W. revised the paper.

