## Supplemental Figure 1 for "Memory-Phenotype Ly49+ CD8 T emerging from the thymus develop into two subsets with distinct immune functions"

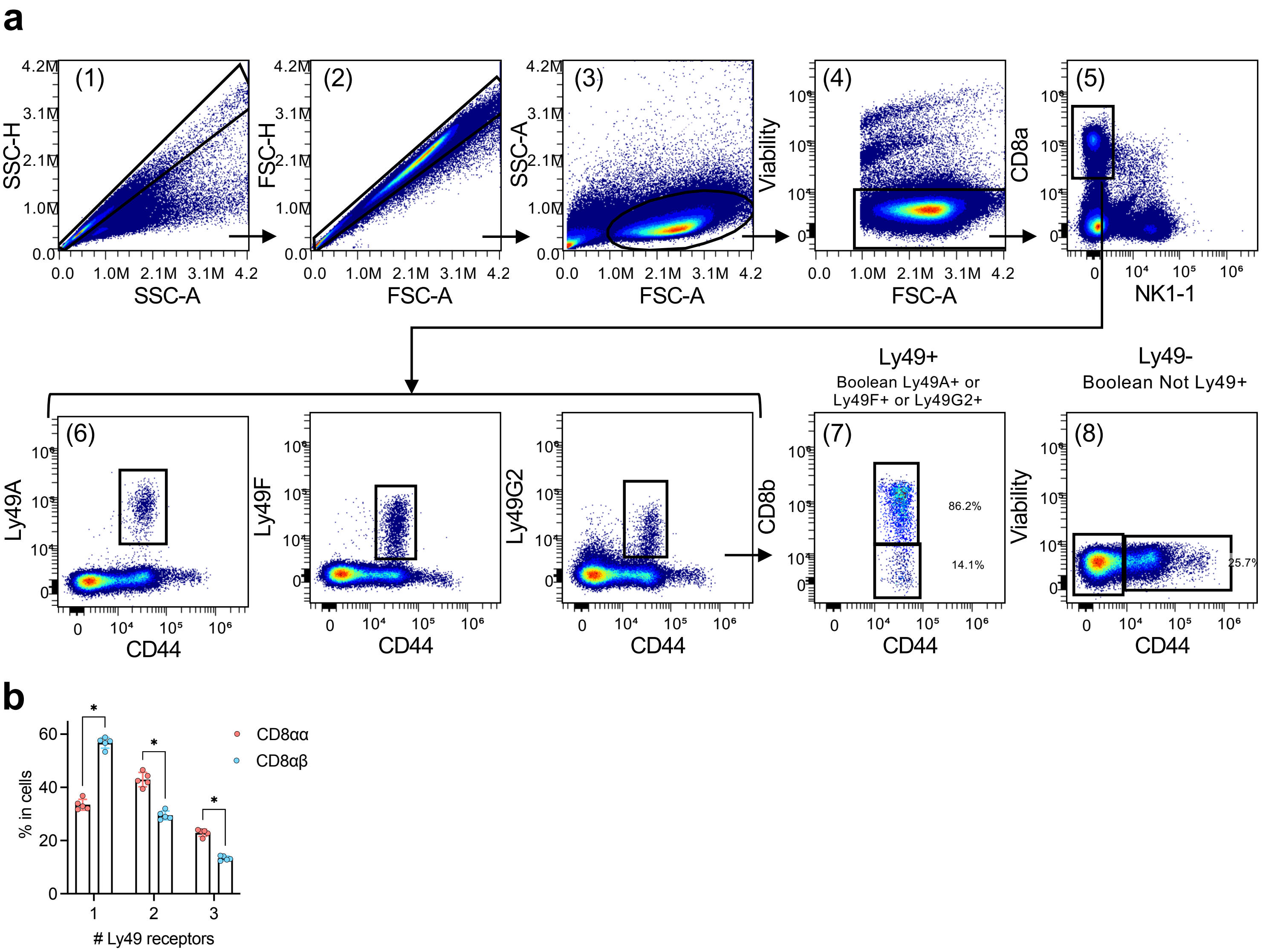


**Supplementary Figure 1 : Characterization of Ly49+ CD8 T cells**

**(a)** Gating strategy of Ly49+ and Ly49- CD8 T cells. (1-2) Doublets cells were eliminated. (3) Lymphocytes were selected and (4) dead cells were eliminated according to viability dye labelling. (5) CD8 T cells were selected and NK were excluded. (6) Ly49+ were selected according to their expression of CD44, Ly49A, Ly49F and Ly49G2. (7) Ly49+ were determined as cells expressing Ly49A, Ly49F or Ly49G2. (8) Ly49- were determined as cells that do not express Ly49A, Ly49F or Ly49G2. (**b**) Proportion of cells expressing 1, 2 or 3 different Ly49 receptors simultaneously with Ly49+ CD8 T cells subsets. Data are represented as mean ± SD (n= 5 mice/group). Statistical significance of differences was determined with Mann-Whitney test (* $p < 0.05$).
