## Supplemental Figure 2 for "Memory-Phenotype Ly49+ CD8 T emerging from the thymus develop into two subsets with distinct immune functions"

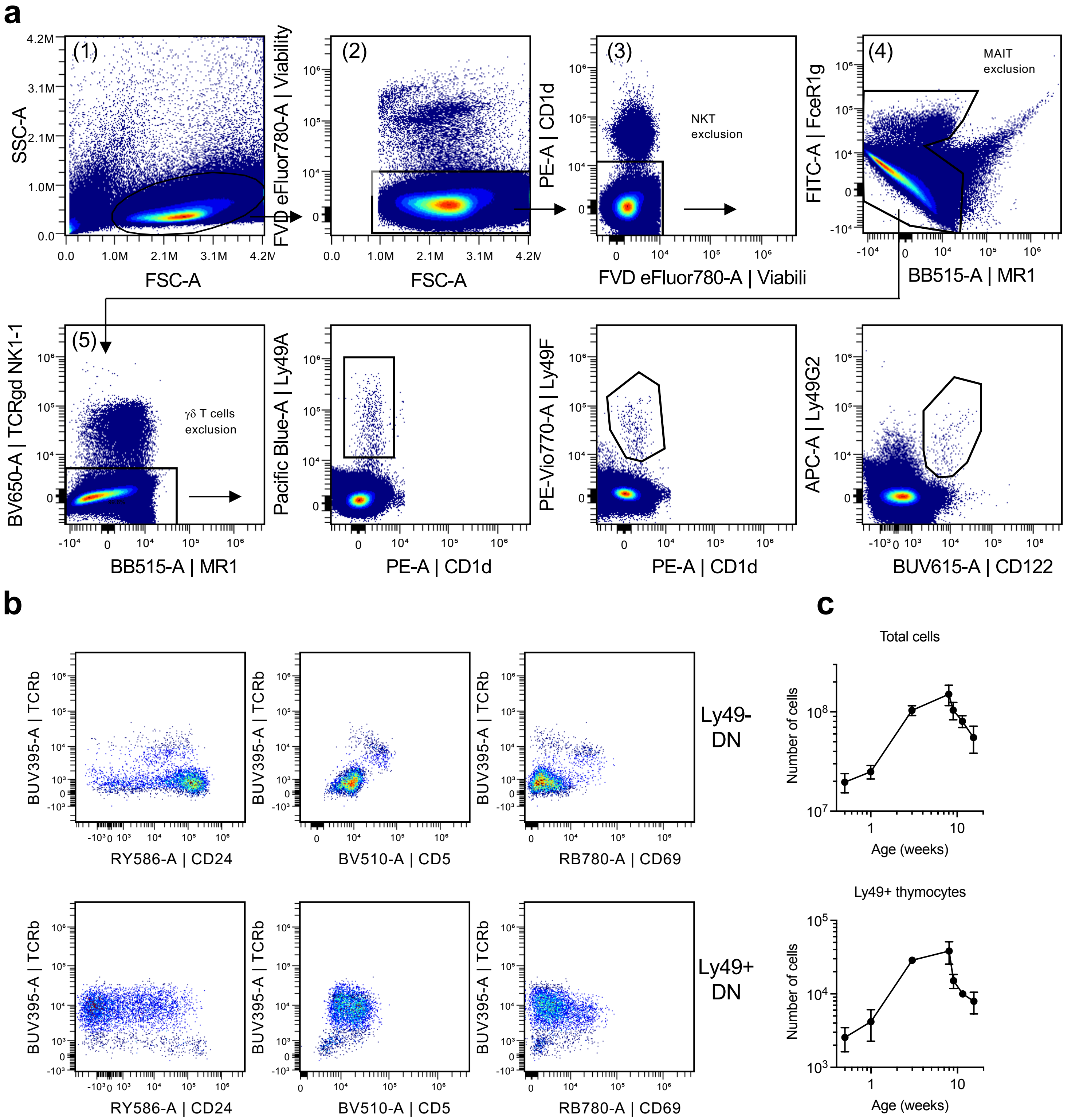


**Supplementary Figure 2 : Characterization of Ly49+ thymocytes**

(**a**) Gating strategy of Ly49+ thymocytes. After exclusion of doublets and selection of lymphocytes (1), viable cells were selected (2) and NKT, MAIT and γδT cells were excluded according to labelling with PB-S57/CD1d tetramer, MR1-5-OP-RU tetramer, anti-NK1.1 and anti-γδ TCR antibody (3-5). Ly49+ thymocytes were then selected according to their expression of Ly49A, Ly49F or Ly49G2 receptors. (**b**) Expression of CD5, CD69 and CD24 by Ly49+ and Ly49- DN thymocytes. (**c**) Number of total cells and Ly49+ cells in the thymus over time.
