## Supplemental Figure 3 for "Memory-Phenotype Ly49+ CD8 T emerging from the thymus develop into two subsets with distinct immune functions"

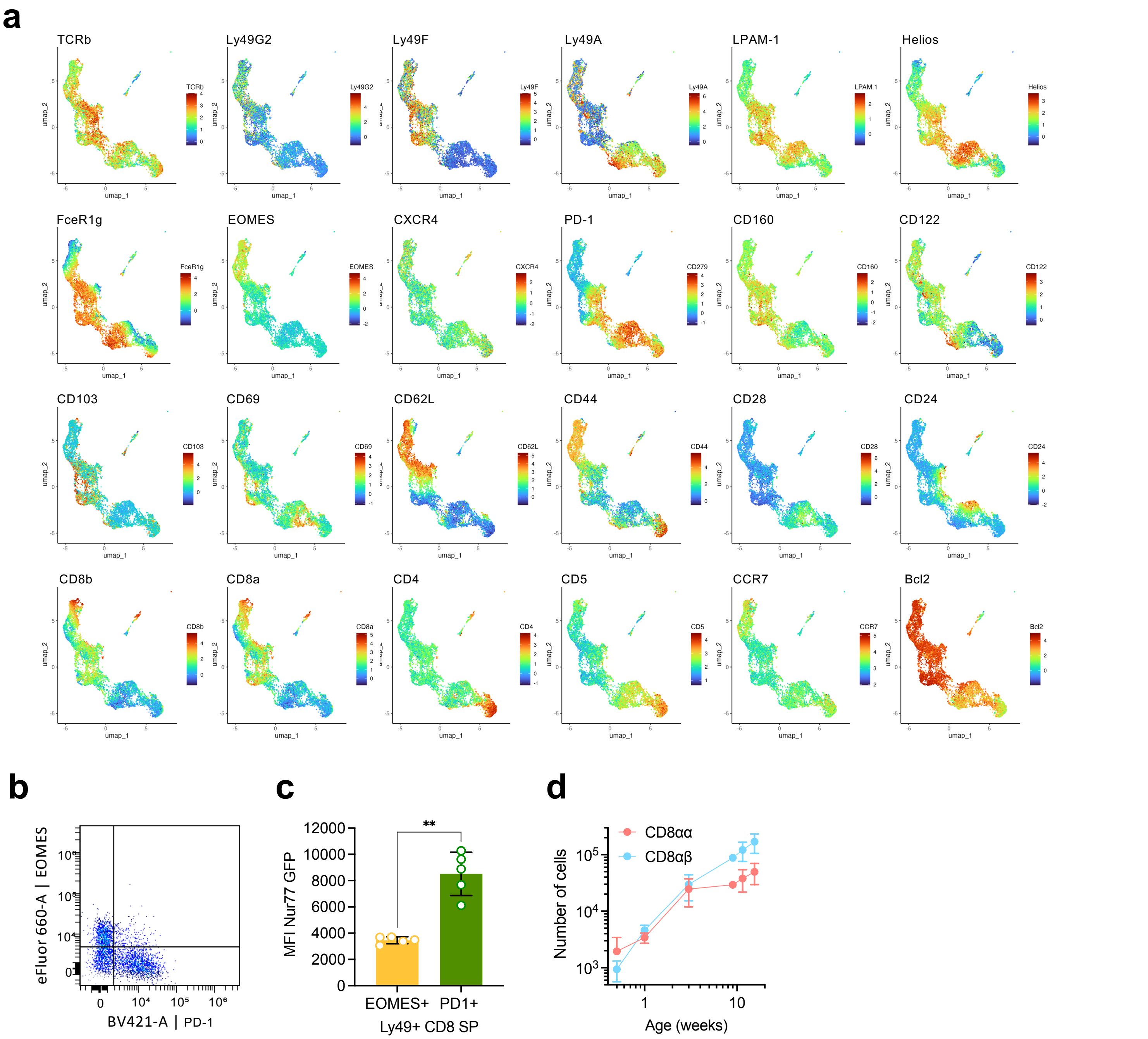


**Supplementary Figure 3 : Comparison of EOMES+ and PD-1+ subsets**

(**a**) UMAP plots showing the expression of the different markers used to determine Slingshot trajectories. (**b**) Dot plots of EOMES versus PD-1 expression within Ly49+ CD8 SP thymocytes. (**c**) Comparison of Nur77-GFP expression between EOMES+ and PD-1+ thymocytes subsets. (**d**) MP-CD8αα-Ly49+ and CD8αβ-Ly49+ cells evolution with age in the spleen. Data are represented as mean ± SD (n = 5 mice/group). Statistical significance of differences was determined with Mann-Whitney test (** p <0.01).
