## Supplemental Figure 4 for "Memory-Phenotype Ly49+ CD8 T emerging from the thymus develop into two subsets with distinct immune functions"

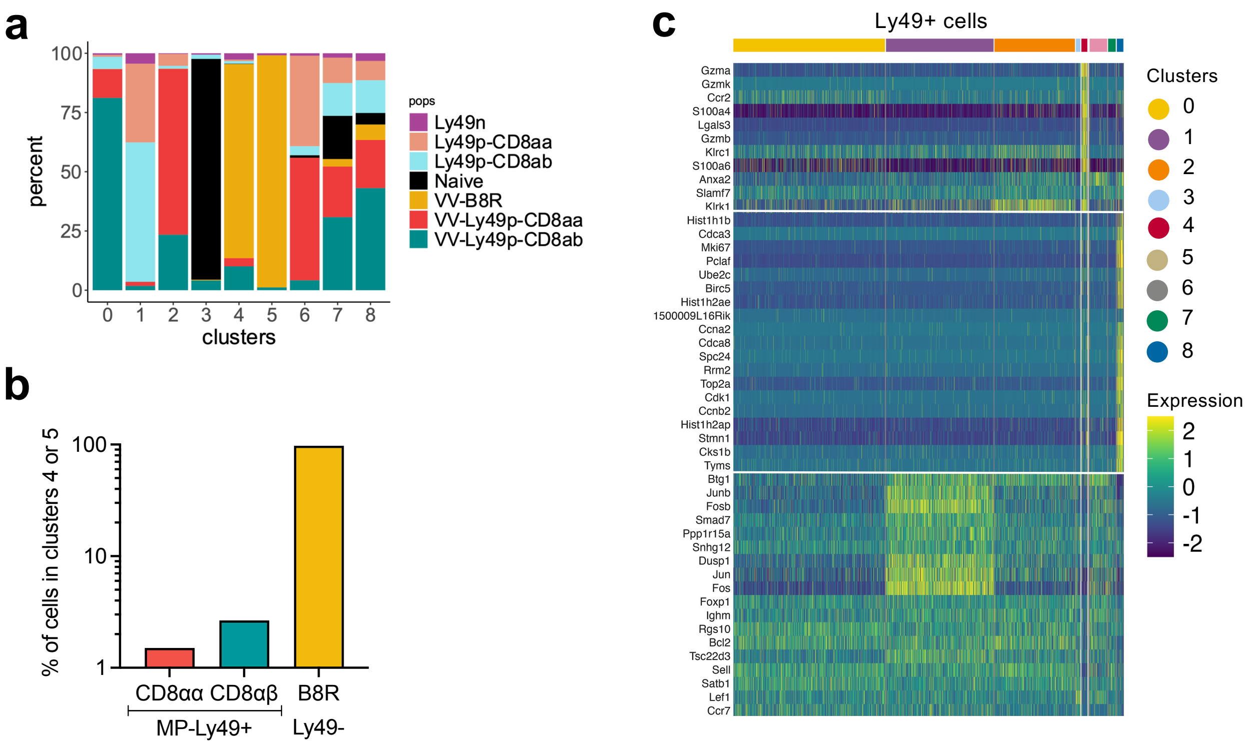


**Supplementary Figure 4 : Transcriptomic analysis of MP-Ly49+ CD8 T cells following viral infection**

(**a**) Proportion of the different cell population sorted as in Fig.4a within each cluster. (**b**) Proportion of MP-CD8αα-Ly49+, MP-CD8αβ-Ly49+ cells and B8R-specific cells from VV-infected mice within clusters 4 or 5. (**c**) Heatmap of the genes differentially expressed in B8R-specific cells (cluster 4 and 5) compared to naive cells (cluster 3), represented for the Ly49+ CD8 T cells clusters.
