## Supplemental Figure 5 for "Memory-Phenotype Ly49+ CD8 T emerging from the thymus develop into two subsets with distinct immune functions"

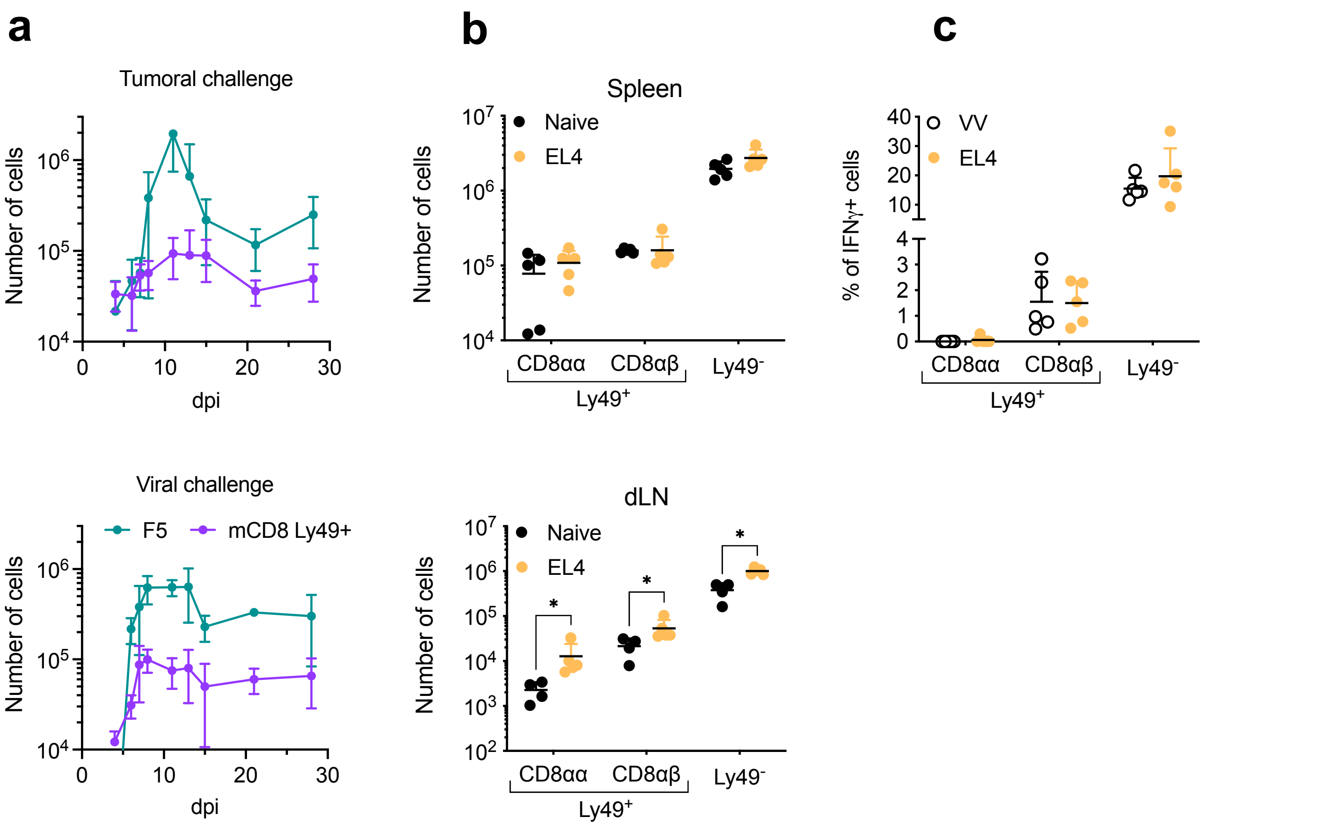


**Supplementary Figure 5 : MP-Ly49+ CD8 T cells can proliferate and respond to a tumoral challenge**

Naive F5 x CD45.1 cells (2.10$^5$) were i.v. transferred in B6 mice 1-day prior immunisation with VV-NP68 (i.n., 2.10$^5$ pfu) or EL4-NP68 cells (s.c., 2,5.10$^6$ cells). (**a**) The number of F5 and MP-Ly49+ CD8 T cells was followed in the blood over time. (**b**) The number of MP-Ly49+ and Ly49- CD8 T cells was measured in the spleen and lymph nodes 8 days post-tumoral challenge. (**c**) B6 mice were immunized with VV-NP68 (i.n., 2.10$^5$ pfu) or EL4-NP68 cells (s.c., 2,5.10$^6$ cells). Height days post-challenge, splenocytes were stimulation with NP68 peptide (10 nM) for 4 hours in the presence of GolgiStop, and the production of IFNγ was measured by flow cytometry. Data are represented as mean ± SD (n = 5 mice/group). Statistical significance of differences was determined with Mann-Whitney test (* $p < 0.05$).
